# Structural covariance networks in children with autism or ADHD

**DOI:** 10.1101/110643

**Authors:** R.A.I. Bethlehem, R. Romero-Garcia, E. Mak, E.T Bullmore, Simon Baron-Cohen

**Affiliations:** Autism Research Centre, Department of Psychiatry, University of Cambridge, Cambridge, CB2 8AH, United Kingdom; Department of Psychiatry, University of Cambridge, Cambridge, CB2 0SZ, UK; Cambridgeshire and Peterborough NHS Foundation Trust, Huntingdon, PE29 3RJ, UK; MRC/Wellcome Trust Behavioural and Clinical Neuroscience Institute, University of Cambridge, Cambridge, CB2 3EB, UK; Academic Discovery Performance Unit, GlaxoSmithKline R&D, Stevenage SG1 2NY, UK; CLASS Clinic, Cambridgeshire and Peterborough NHS Foundation Trust, United Kingdom

**Author notes:** Corresponding author: Richard A.I. Bethlehem, Autism Research Centre, Douglas House, 18b Trumpington Road, CB2 8AH, Cambridge, UK. Authors contributed equally (R.A.I. Bethlehem, R. Romero-Garcia).

**Keywords:** structural covariance, autism, ADHD, cortical thickness, graph theory

## Abstract

While autism and attention-deficit/hyperactivity disorder (ADHD) are considered distinct conditions from a diagnostic perspective, they share some phenotypic features and have high comorbidity. Taking a dual-condition approach might help elucidate shared and distinct neural characteristics.

Graph theory was used to analyse properties of cortical thickness structural covariance networks across both conditions and relative to a neurotypical (NT; n=87) group using data from the ABIDE (autism; n=62) and ADHD-200 datasets (ADHD; n=69). This was analysed in a theoretical framework examining potential differences in long and short range connectivity.

We found convergence between autism and ADHD, where both conditions show an overall decrease in CT covariance with increased Euclidean distance compared to a neurotypical population. The two conditions also show divergence: less modular overlap between the two conditions than there is between each condition and the neurotypical group. Lastly, the ADHD group also showed reduced wiring costs compared to the autism groups.

Our results indicate a need for taking an integrated approach when considering highly comorbid conditions such as autism and ADHD. Both groups show a distance-covariance relation that more strongly favours short-range over long-range. Thus, on some network features the groups seem to converge, yet on others there is divergence.

## Introduction

Autism spectrum conditions (henceforth autism) are characterized by deficits in social communication alongside unusually restricted interests and repetitive behaviours, difficulties adjusting to unexpected change, and sensory hypersensitivity (American Psychiatric Association 2013). Despite a large body of research to understand its underlying neurobiology (Loth et al. 2015), no distinct set of biomarkers for autism has yet been established. With respect to the neuroimaging literature and specifically network connectivity, several authors have suggested potential differences in brain organization in autism compared to neurotypical control groups. There is however debate about whether autism is characterized by neural over-or under-connectivity (Brock et al. 2002; Rubenstein and Merzenich 2003; Belmonte et al. 2004; Just et al. 2004; Courchesne and Pierce 2005). The working hypothesis is that people with autism suffer from atypical connectivity (Courchesne and Pierce 2005; Cherkassky et al. 2006; Just et al. 2007; Assaf et al. 2010). Specifically, there is a tendency for autism to be associated with excess local or short-range connectivity, relating to enhanced local processing. This is thought to be accompanied by decreased global or long-range connectivity, relating to impaired integration as manifested in ‘weak central coherence’. Thus, a prominent theory of neural connectivity in autism is of global under- and local over-connectivity (Belmonte et al. 2004; Vissers et al. 2012). Attention-deficit/hyperactivity disorder (ADHD) on the other hand is characterised by a triad of symptoms: hyperactivity, impulsive behaviour and inattentiveness (American Psychiatric Association 2013). Studies using connectivity analyses have attempted to shed light on its underlying neurobiology and have found both decreased and increased functional connectivity in specific networks (Tomasi and Volkow 2012), altered connectivity in the default mode network (DMN) (Fair et al. 2010) and differences in cross-network interactions (Cai et al. 2015). These effects might be smaller than literature suggests (Mostert et al. 2016).

Autism and ADHD show high comorbidity and phenotypic overlap (Rommelse et al. 2010, 2011; Leitner 2014), and are both also potentially marked by differences in connectivity. There have even been suggestions that these connectivity differences lie on a similar dimension of local and global connectivity imbalances (Kern et al. 2015). In addition, both conditions have been associated with alterations in cortical development (Shaw et al. 2007; Hardan et al. 2009) that could in turn give rise to differences in the topological organisation of brain networks. In the present study, we aimed to identify distinct as well as overlapping patterns of brain organisation that might shed a light on the underlying architecture of both conditions, giving rise to divergent yet related findings using structural covariance analyses.

Structural covariance analysis involves covarying inter-individual differences (i.e. coordinated variations in grey matter morphology) in neural anatomy across groups (Alexander-Bloch *et al.,*2013; Evans, 2013) and is emerging as an efficient approach for assessing structural brain organization. A key assumption underlying this methodology is that morphological correlations are related to axonal connectivity between brain regions, with shared trophic, genetic, and neurodevelopmental influences (Alexander-Bloch *et al.,* 2013). Thus, structural covariance network analysis is not the same as analysis of functional connectivity or structural networks obtained with diffusion imaging, yet it has shown moderately strong overlap with both (Alexander-Bloch *et al.,* 2013; Gong *et al.,* 2012). In addition, structural covariance networks are highly heritable (Schmitt et al. 2009) and follow a pattern of coordinated maturation (Alexander-Bloch *et al.,* 2013; Raznahan *et al.,* 2011; Zielinski *et al.,* 2010). With respect to neurodevelopmental conditions, structural covariance networks might provide a way to investigate potential differences in brain network development. Differences between neurotypical individuals and individuals with a developmental condition are likely the result of divergent developmental trajectories in coordinated development of different brain networks. The advantage of structural covariance analysis is that it focuses on this coordinated structure of the entire brain as opposed to focusing on a specific structure. In addition, structural data on which these networks are based is widely available, analysis is less computationally intensive and arguably less sensitive to noise, compared to functional imaging.

Previous investigations of structural covariance in autism have shown regional or nodal decrease in centrality, particularly in key regions subserving social and sensorimotor processing, compared to neurotypical individuals (Balardin et al. 2015). Furthermore, speech and language impairments in autism have been associated with differences in structural covariance properties (Sharda et al. 2014). Studies of functional connectivity networks in autism are more abundant (Vissers et al. 2012). In ADHD structural covariance analyses have been extremely scarce. A study that specifically investigated structural covariance in drug-naïve adolescent males found that grey matter volume covariance was significantly reduced between multiple brain regions including: insula and right hippocampus, and between the orbito-frontal cortices (OFC) and bilateral caudate (Li et al. 2015). Similar to the autism literature, studies that have explored functional connectivity differences in ADHD are more abundant (Konrad and Eickhoff 2010).

While autism and ADHD are considered distinct conditions from a diagnostic perspective, clinically they share some common phenotypic features (such as social difficulties, atypical attentional patterns, and executive dysfunction) and have high comorbidity (Rommelse et al. 2010, 2011; Leitner 2014). DSM-5 (American Psychiatric Association 2013) now allows comorbid diagnosis of autism and ADHD, acknowledging the common co-occurrence of these conditions. Regardless, most studies to date have focused on each condition separately, with considerable heterogeneity in results. Taking a dual-condition approach might help elucidate shared and distinct neural characteristics. Our proposal for a dual-condition approach is supported by a recent review that found both distinct as well as overlapping neural characteristics between autism and ADHD (Dougherty et al. 2015). There is also increasing interest in the clinical and research communities to investigate autism and ADHD along a continuum of atypical neural connectivity (Kern et al. 2015).

In the present study, we used the graph theoretical framework to analyse properties of structural covariance networks across autism and ADHD, relative to an age and gender matched neurotypical control (NT) group. One study has taken a similar approach using resting-state fMRI and diffusion weighted tractography and reported marked connectivity differences between network hubs, indicating a disruption in rich-club topology. Specifically, Ray and colleagues (Ray et al. 2014)report a decrease in connectivity within the rich-club but increased connectivity outside the rich-club in ADHD. The autism group showed an opposite pattern of increased connectivity within rich-club connectivity. These findings may fit with the idea of increased local connectivity in autism (i.e., increased within rich-club connectivity), with ADHD showing the opposite pattern. Yet, these findings could also mediate increased strength in long-range connections within the rich-club. In the present study we aimed to further investigate the relation between distance and connectivity by looking at group-wise cortical thickness covariance as a function of Euclidean distance. In addition, we investigate potential overlap in modular and hub organization as assessed by structural covariance network analyses.

## Methods

### Image processing and quality control

Structural T1-weighted MPRAGE images were collected from two publically available datasets: ABIDE (http://fcon_1000.projects.nitrc.org/indi/abide/) and ADHD-200 (http://fcon_1000.projects.nitrc.org/indi/adhd200/). From these datasets, 3 diagnostic groups (autism, ADHD and neurotypical individuals) of males between the ages of 8 and 12 years old were selected. The initial sample consisted of 348 eligible individuals. The structural T1-MPRAGE data were pre-processed using Freesurfer *v5.3* to estimate regional cortical thickness. Cortical reconstructions were checked by three experienced independent researchers. Images were included in the analyses only when a consensus on the data quality was reached (see Supplementary Information for more details on data selection). The cortical thickness maps were automatically parcellated into 308 equally sized cortical regions of 500 mm^2^ that were constrained by the anatomical boundaries defined in the Desikan-Killiany atlas (Desikan et al. 2006; Romero-garcia et al. 2012). The backtracking algorithm grows subparcels by placing seeds at random peripheral locations of the standard atlas regions and joining them up until a standard pre-determined subparcel size is reached (Romero-garcia et al. 2012). It does this reiteratively (i.e., it restarts at new random positions if it fails to cover an entire atlas region) until the entire atlas region is covered. Individual parcellation templates were created by warping this standard template containing 308 cortical regions to each individual MPRAGE image in native space. A key advantage of warping of the segmentation map to the native space relates to the attenuation of possible distortions from warping images to a standard space that is normally needed for group comparisons. Lastly, average cortical thickness was extracted for each of the 308 cortical regions in each individual participant.

As a secondary post-hoc step in quality control, individuals that had an average variability in cortical thickness of more than two standard deviations away from the group mean were removed from further analysis. After quality control and matching on age and IQ, our final sample consisted of 218 participants: ADHD (*n=69, age = 9.99 ±1.17, IQ = 107.95 ±14.18),* autism (*n=62 age=10.07 ±1.11, IQ = 108.86 ±16.94)* and NT (*n=87, age = 10.04 ±1.13, IQ = 110.89 ±10.39*). See S1 Fig. S1 for an overview and Table S1 for details on scanner site and matching procedure. Scanner site was regressed out from raw cortical thickness estimates across groups. To aid interpretation of the cortical thickness estimates, the residuals from this regression where added to the sample mean. Group-wise structural covariance matrices were then computed by taking the inter-regional Pearson correlation of these parcel-wise cortical thickness estimation. This was done within each group to create group-wise structural covariance matrices.

## Data Analysis

### Distance connectivity differences

To determine potential group effects on the CT covariance for short and long range associations, we investigated the linear slope differences in the relationship between correlation strength and Euclidean distance between nodal centroids. Consequently, oneway analysis of covariance (ANCOVA) were performed with the diagnosis group as a factor and Euclidean inter-regional distance as a covariate. For significant group effects, post-hoc paired t-tests were used to identify which slopes are significantly different from each other.

### Graphs

To construct adjacency matrices for graph analyses, the minimal spanning tree (van Wijk et al. 2010) was used as the threshold starting point for building covariance networks at a representative density of 10%. The density of a network relates to the fraction of edges present in the network compared to the maximum possible number of edges. Graph analyses were performed across densities and between-group differences were compared using non-parametric permutation tests on paired group comparisons (1000 permutations). Thus, permuted networks were constructed by permuting the underlying cortical thickness estimates for each group comparison and constructing adjacency matrices for each. In view of the large number of comparisons across the 308 nodes, differences in local measures were subjected to a False Discover Rate (FDR) non-linear multiple comparison correction with alpha set at < 0.025 to allow simultaneous correction for two-tailed testing (Benjamini and Hochberg 1995).

### Degree, cortical thickness and wiring cost analysis

Nodal degree reflects the number of edges connecting each node. Nodes with the highest degree of the network are defined as hubs. The present study considered a wide range of degree thresholds to reduce bias related to the choice of an arbitrary set of hubs (ranging from 0 to 100% of the nodes). Thus, group differences in degree and CT of the hubs of the networks were evaluated for each degree threshold. To decrease the noise effect, we calculated the cumulative degree distribution as *P*(*k*)= Σ_*k*′≥*k*_ p(*k*′).

Inter-regional distance (*d_ij_*) between two nodes *i* and *j* was estimated as the Euclidean distance between the centroids, 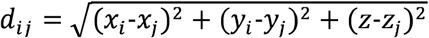, where *x*, y and *z* represents the coordinates of the centroid of each region in MNI space. The mean connection distance or wiring cost (*W_c_*) of a network was computed as, *W_c_ =* (*Z_iij_net_ij_^*^d_ij_*)*/N,* where *net*(*i,j*) is equal to 1 if regions *i* and *j* are connected, 0 otherwise and *N* is the total number of connections of the network.

### Modular agreement

Modular agreement was evaluated by quantifying the proportion of pairs of regions that were classified within the same module in community partitions (using iterating Louvain clustering to obtain modular partitions) associated with different diagnostic groups. Thus, two groups will show high modular agreement if network modules mainly include the same set of brain regions in both groups. As modular agreement is highly affected by intrinsic trivial characteristics of the modular partition, z-scores were used as a measure of how over- or under-represented a given metric was compared with random community partitions. In order to test against appropriately designed surrogate data, statistical significance was assessed against a null distribution built from metric values computed in 1000 random communities generated by preserving the number of modules, size of the modules, spatial contiguity and hemispheric symmetry of the real community partition. The 95^th^ quantile of the resulting distribution was used as a statistical threshold to retain or reject the null hypothesis of no significant modular agreement between diagnostic groups. Moreover, differences in modular agreement between pairs of groups were statistically tested using a similar procedure. Indices of modular agreement of each pair of groups were subtracted and compared with the differences of modular agreement derived from the 1000 random communities in each pair of groups. Similarly, the 95^th^ quantile of the resulting distribution was used as a statistical threshold to retain or reject the null hypothesis of no modular agreement differences between pairs of diagnostic groups. Significant results were corrected for multiple comparisons using FDR (Benjamini and Hochberg 1995).

## Results

### Distance covariance topology

In all groups the group-wise correlation strength decreased with increased anatomical distance. Results from the analysis of variance show a main effect of group F(2,141828) = 2192.76, p<0.0001. Post-hoc analyses indicated that all three group have a small but significantly different slope: ADHD < Neurotypical (p-value < 10^−15^), Autism < Neurotypical (p-value < 0.005) and ADHD < Autism (p-value < 10^−15^). Figure 1 shows the linear relation of the inter-regional correlation as a function of Euclidean distance and the mean and confidence intervals of the slope estimates. In the ADHD group, inter regional correlation decreased the fastest whereas the neurotypical group shows the smallest decreases. This result shows that both autism and ADHD have relatively weaker long-range covariance and stronger local covariance. Compared to the neurotypical group both groups show a balance that more strongly favors short-range over long-range covariance.

**Figure 1:**
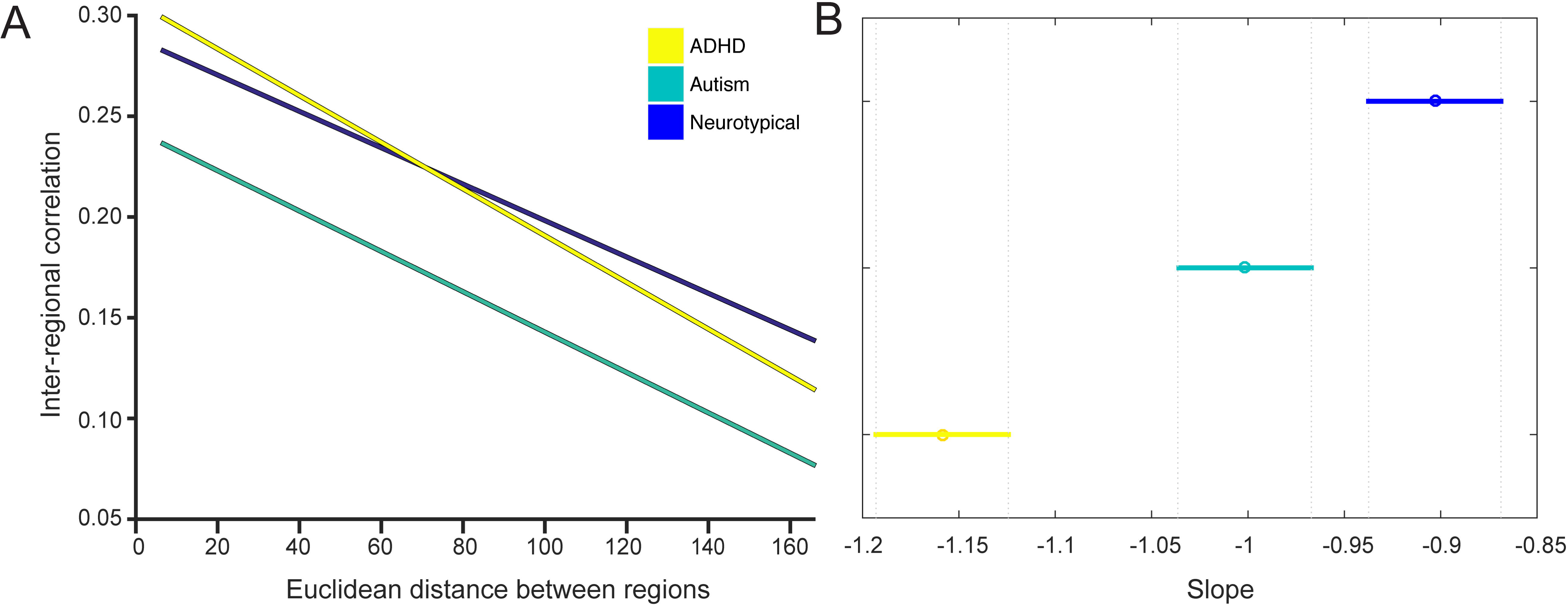
Inter-regional correlation strength as a function of Euclidean distance. Panel A shows the inter-regional correlation over the entire distance range. Panel B shows the mean slope for each group and the 95% confidence interval of the mean slope.

### Degree

After constructing the covariance matrices (Figure 2A), the degree of each node was computed (Fig. 2B) and the top 10% nodes with highest degree were retained as hubs for visualization (Fig. 2C). Most of the hubs were located within frontal and parietal cortices in the three groups. On the contrary, nodes with lower degree were mainly placed in the occipital cortex. There were several nodes that showed degree differences between groups, but these were not consistent across degree densities. We did however observe marked differences between groups in the overall degree distribution. Figure 3 show the cumulative degree distribution of each group. Interestingly, hubs of the autism group exhibited significantly lower degree than both neurotypical (p-value < 0.025; for degree values from 83 to 88) and ADHD (p-value < 0.025; for degree values from 64 to 89). These difference were corrected for multiple comparisons for the range of higher degree nodes (FWE correction in the degree range from 50 to 90).

**Figure 2:**
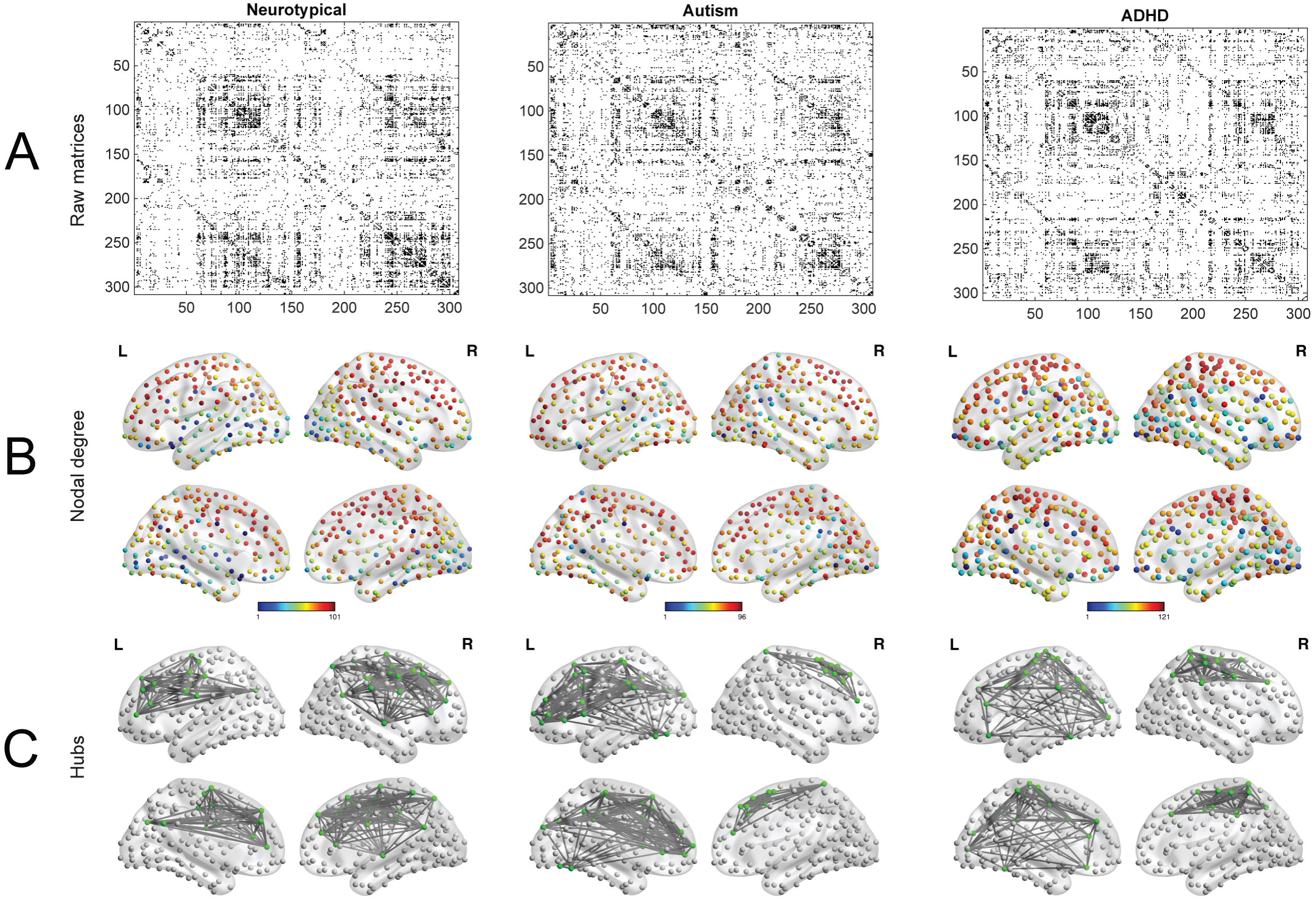
Overview of procedure and metrics. **Panel A** shows the binary adjacency matrices for the three groups thresholded at **10%** above the minimal spanning tree. Subseguent graph construction is based on these thresholded matrices. **Panel B** display the topological distribution of nodal degree at **10%** density. **Panel C** illustrates the networks with nodes that have the highest degree (top 10%).

**Figure 3:**
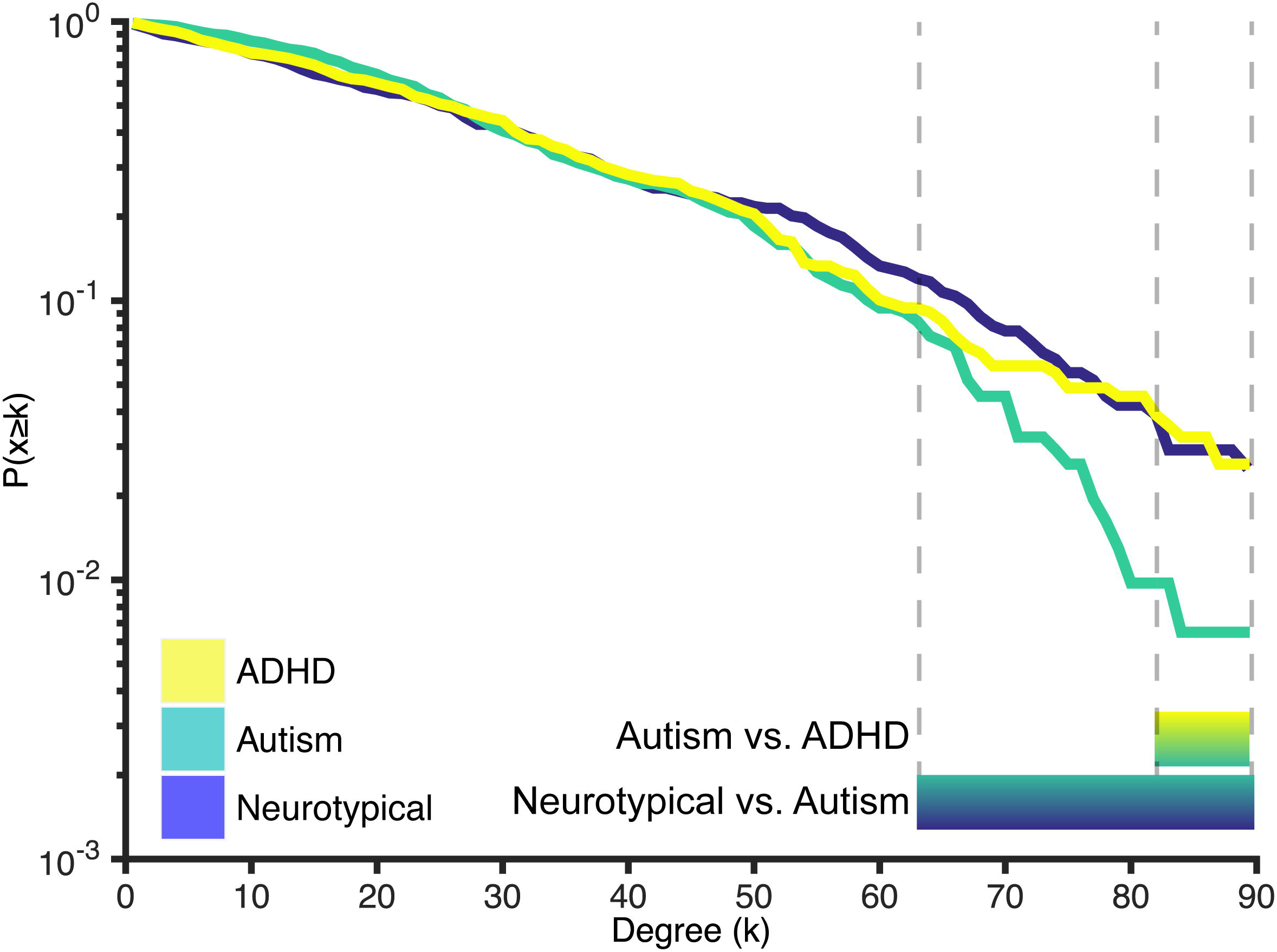
Cumulative degree distribution. Lines represent the proportion of nodes in the network with a degree higher than k (hubs) in each group. Bars below the figure represent the areas where there is a significant difference between the groups. Hubs of the autism group showed significantly lower degree compared to the ADHD group (k-range: 83-88) and compared to the neurotypical group (k-range: 64-89).

### Wiring cost

In line with the group differences observed in the decay of cortical thickness correlation as a function of the inter-regional distance described above, the wiring cost analysis showed a significant decrease of the average distance between connected regions in the ADHD group compared with neurotypical (Figure 4; p-value<0.008), revealing a reduction of long range connections in the ADHD network.

**Figure 4:**
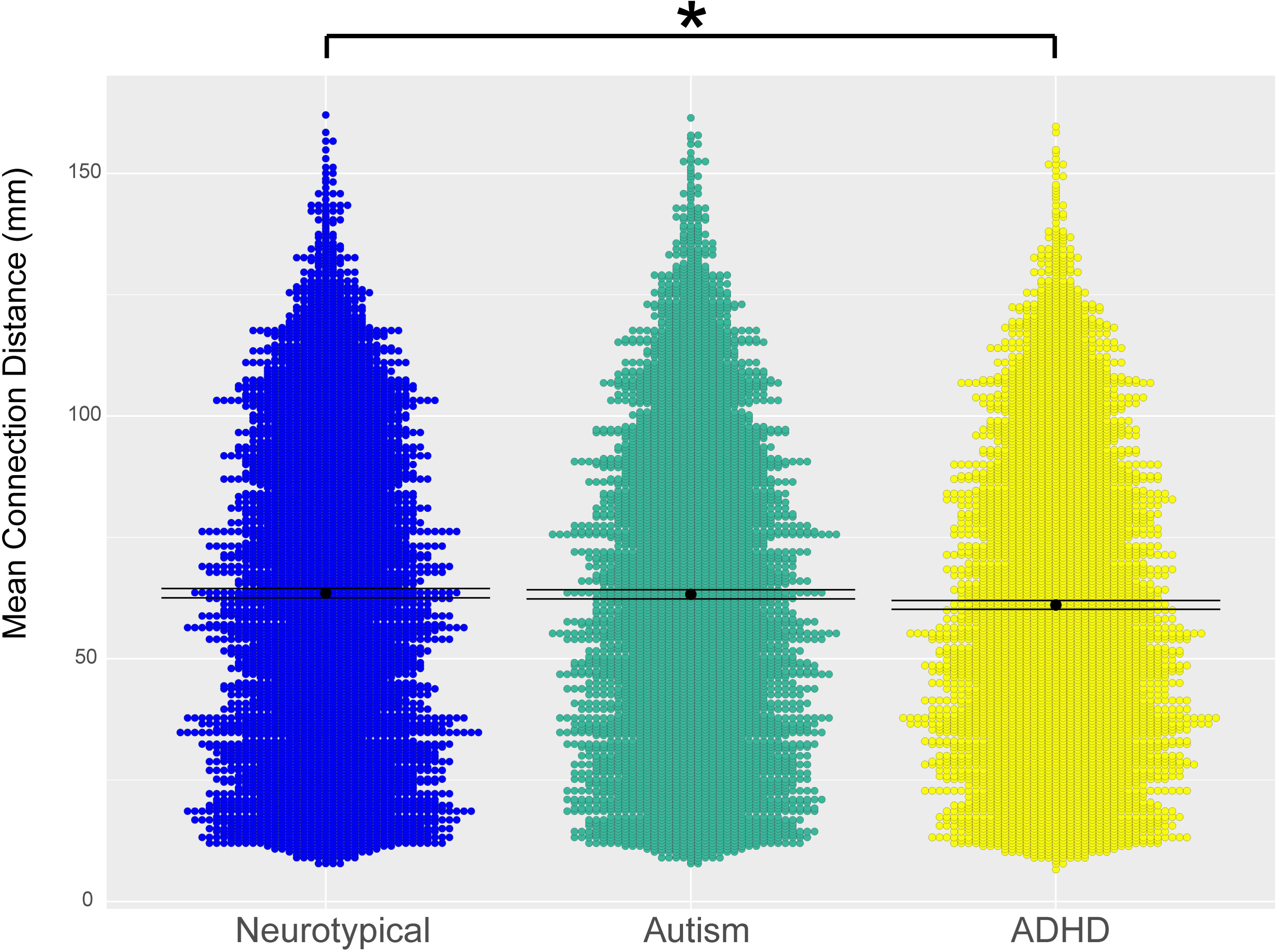
Violin representation of the mean inter-regional distance between connected regions in the three groups. The ADHD group has significantly lower connection distance compared to the neurotypical group. Mean is shown as a black dot with error bars representing 95% confidence intervals

### Cortical thickness as a function of degree

Given that there were notable differences in degree distributions (i.e. hubs in the autism group had lower degree than the other groups; Figure 2) we chose to analyze both the absolute degree distribution and take a percentile that was based on the group itself. Although the autism and neurotypical group showed little difference in cortical thickness across the entire range of degrees with both methods, high degree nodes had significantly reduced cortical thickness in the ADHD group (Fig. 5). This suggests that there might be increased synaptic pruning in these hub regions in the ADHD group.

**Figure 5:**
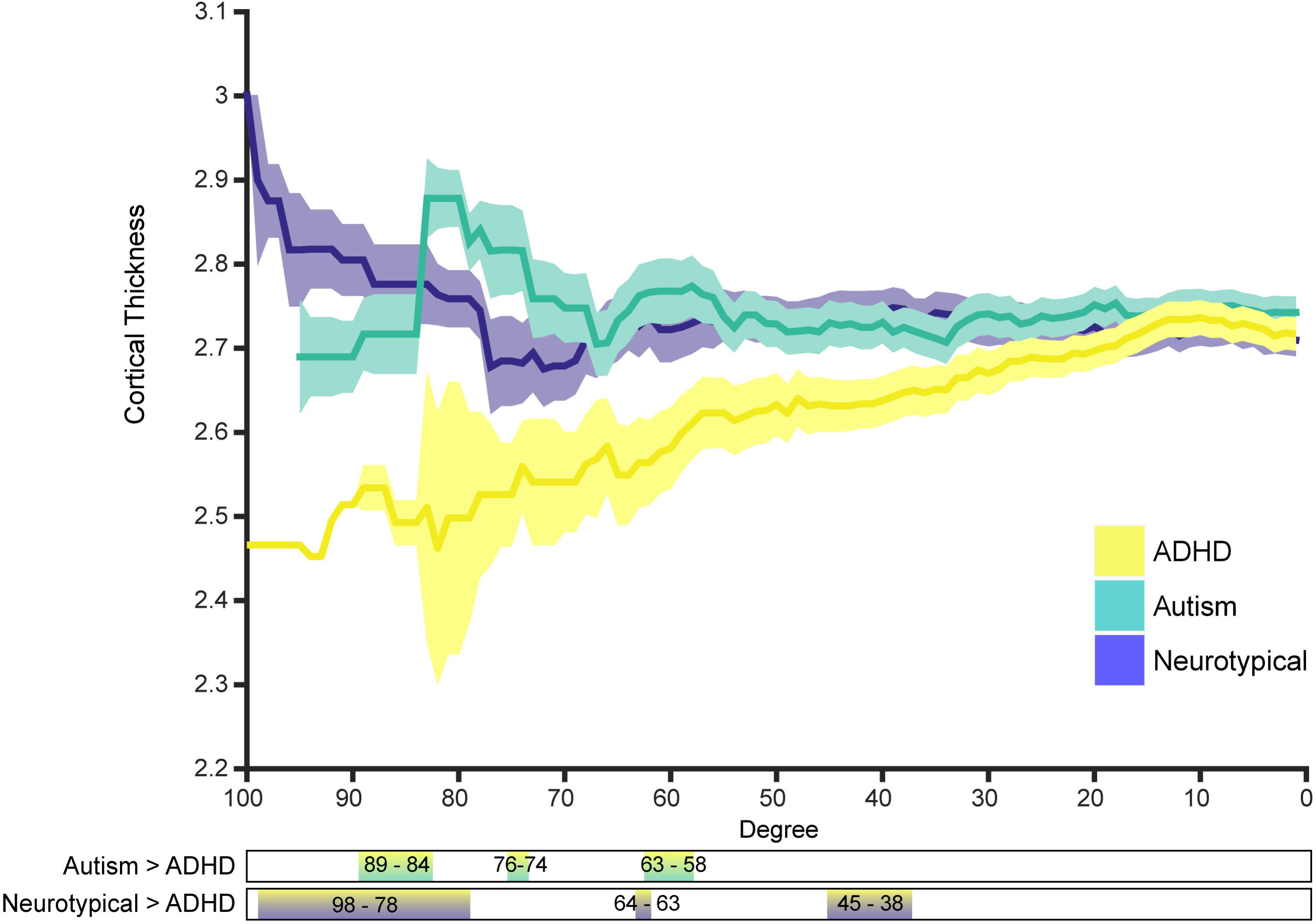
Cortical thickness as a function of degree. Bars below the figure show the degree ranges where there is a significant difference between the respective groups.

### Modular consistency and clustering

To investigate similarities in global topology, we further evaluated the modular overlap between the community structure of the three groups. The modular overlap between all group-wise comparisons were significantly higher than expected by chance (Fig. 6), suggesting that a global scale there were no marked differences in structural covariance community structure. However, the Autism-ADHD group overlap was significantly lower than the Neurotypical-ADHD overlap (p-value<10^−3^). There was also a small non-significant effect for the Neurotypical-ADHD overlap compared to Neurotypical-Autism (p-value=0.04). This indicates that although there were perhaps no massive topological differences in community structure, the autism and ADHD group differ more from one another than they do from the neurotypical group (i.e. there was lower modular agreement between autism and ADHD then there was between the other groups).

**Figure 6:**
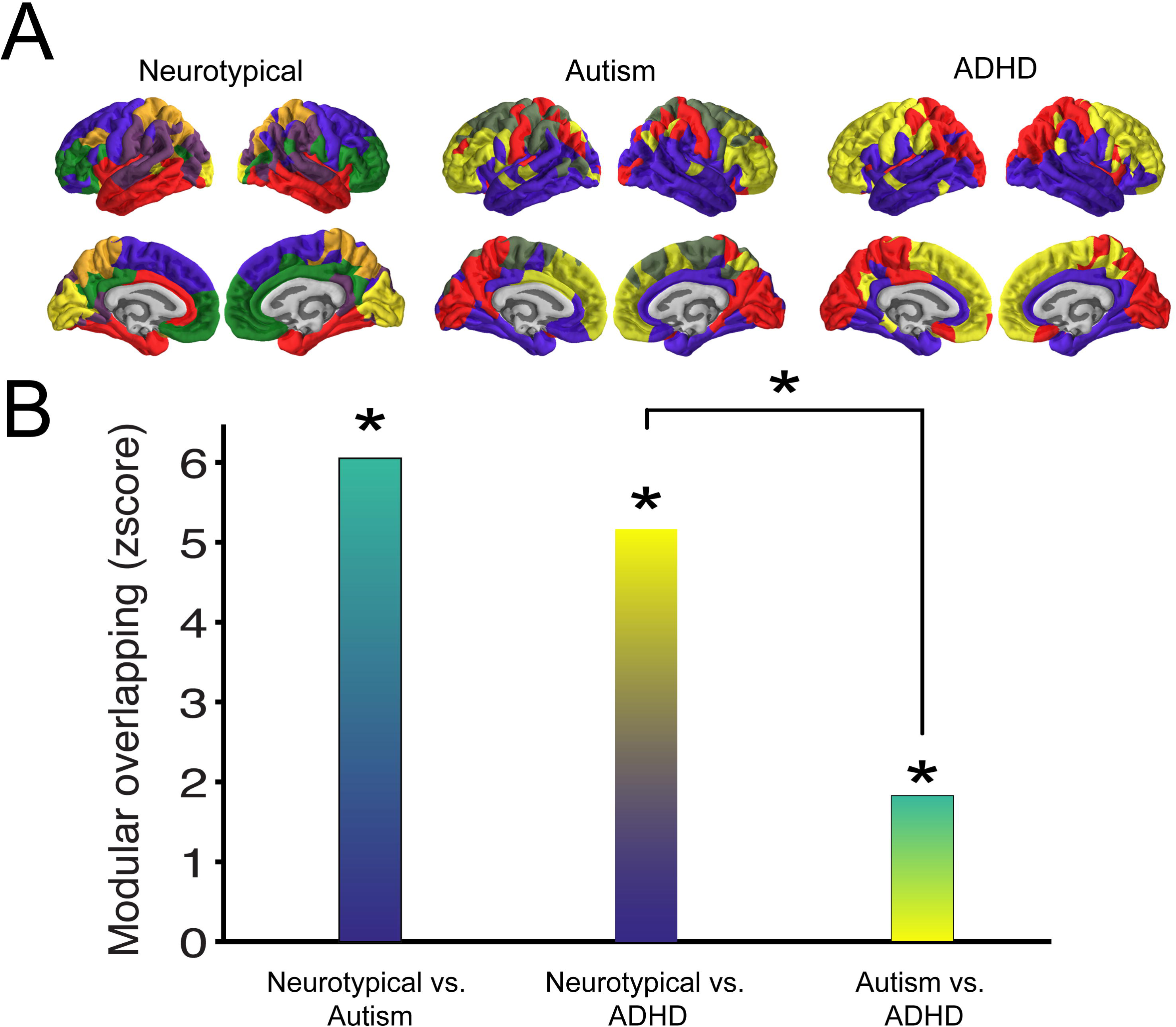
Similarities in community structure across groups. *Panel* ***A*** *illustrates the modular organization of the structural covariance network derived from each group. The colours show association of the region with a certain module. These colours are set for each group individually as not all groups have the same number of modules. Panel* ***B*** *displays the z-transformed modular overlap for each group-wise comparison, colour meshes are chosen to represent the group comparison. All overlap scores are significantly different from zero, indicating that nodes in one module are most likely part of the same module in both groups. Note that Autism-ADHD overlap was reduced compared to the NT-ADHD overlap.*

## Discussion

Comparing autism and ADHD, our findings reveal a complex topology of convergent yet distinct patterns of brain network organization. At a global level of community structure all groups show a significant degree of overlap, however the autism and ADHD group showed less similarity than they do compared to the neurotypical control group. The decay of cortical thickness correlation strength as a function of inter-regional distance was also markedly different for both clinical groups. Fitting with the idea of a local vs global connectivity difference in developmental conditions both the autism and ADHD group showed a pattern that diverges from the neurotypical control group. Yet, they do not appear to be in opposing direction. Both group showed a significantly stronger decrease in correlation strength with increased distance relative to a control group.

These finding seem to suggest that in both conditions the topology favour short-range correlations over long-range correlations. This idea is prominent in autism literature, but less so in the ADHD literature. It will be interesting for future studies on different modalities such as resting-state or DTI imaging to see if potential connectivity differences follow a pattern similar to the present structural covariance properties. In addition, we found that the ADHD group a marked decrease in cortical thickness in high degree regions compared to the other two groups. A previous study showed that children with ADHD exhibited reduced CT in fronto-parietal regions, but increased CT in occipital regions (Almeida Montes et al. 2013). In the present analysis cortical hubs were mainly located in fronto-parietal networks, thus this finding fits with the idea of overall reduced CT in those areas. Interestingly, Almeida-Montes and colleagues also show that some of these difference increase with age. This would also fit with previous work showing some delay in cortical maturation of cerebrum and specifically prefrontal cortex in children with ADHD (Shaw et al. 2007).

A previous study indicated that wiring costs in autism might also fit in a model of increased local connectivity and decreased global connectivity in grey matter connections (Ecker, Ronan, et al. 2013). Thus, we extended our local versus global analysis to include wiring cost characteristics. We found that the ADHD group showed significantly reduced wiring cost. This would be consistent with the notion of a network shift towards increased segregation (i.e. more local connections) at the expense of global integration. We did not find a significant difference in the wiring cost for the autism group. The present approach to assess wiring costs differs significantly from the one taken by Ecker et al (Ecker, Ronan, et al. 2013)(e.g., we use Euclidean distance between centroids of anatomically derived nodes compared to a measure of mean separation distance on the cortical sheet). It is possible that our approach might be too coarse to pick up wiring cost differences in the autism group. Our results do indicate a sharp reduction in the number of connections of the hubs regions in the autism network that can be explained in terms of a shift away from global integration. Again, future studies will have to show whether these patterns also emerge from connectomic data.

Since changes in structural covariance are postulated to be a result of a prolonged developmental process, our findings also provide emerging evidence for a systematic difference in the developmental trajectory/profile of brain organization between these groups. However, a recent large cross condition analysis of potential genetic relationship showed only moderate genetic overlap between autism and ADHD (Lee et al. 2013). Thus, the true underlying cause for these differences is likely more indirect and could emerge from long-term differences in functional connectivity.

Contrary to our predictions, and in contrast to a previous study (Ray et al. 2014) that used a different imaging modality, we did not find any significant differences in rich-club topology between any of the groups. The rich-club coefficient indicates that high degree nodes are more likely to connect to other high-degree nodes (sometimes summarised as ‘the rich cling together’). Although the structural covariance networks were constructed from T1-MPRAGE data, we had expected to find overlap between the fMRI, DTI and our current results. It would be highly interesting to see how these differences develop further. Connectivity findings in adult autism and ADHD are notoriously heterogeneous (Konrad and Eickhoff 2010; Vissers et al. 2012), so some developmental neuroanatomical differences might gradually change with age. The present data was restricted to a very specific age group and developmental changes continue long after this time frame. It would be interesting to see whether the currently observed lack of differences in structural covariance topology propagate in the same direction. More research is needed to assess these potential longitudinal changes in this population.

Modular organization of the network of the three groups revealed no significant differences, but instead showed significant overlap. Therefore, network nodes belonging to one module in one group are likely to belong to the same module in the other group. Considered in the clinical context of overlapping phenotypes and high comorbidity, the present results strengthen the notion that these two conditions should not be studied in isolation. However, the two clinical groups (despite being significantly similar) show less modular similarity to one another than they do compared to a neurotypical group. However, both groups also showed significant overlap with the neurotypical group, suggesting that the neuroanatomical differences between the clinical and control groups operate on more fine-grained scales (such as might be observed in graph theoretical measures). This finding shows that when these groups are studied solely in contrast with a neurotypical group no difference might be observed on this metric.

There are some caveats surrounding the current study. First, and in contrast to some studies, we used cortical thickness estimates to construct our structural covariance network, thereby excluding sub-cortical regions from network analysis. For this reason, some studies have used covariations of grey matter volume instead (Balardin et al. 2015). However, grey matter volume relies on the relationship between two different morphometric parameters, cortical thickness and surface area. Cortical thickness and surface area are both highly heritable but are unrelated genetically (Panizzon et al. 2009), leading to different developmental trajectories across childhood and adolescence (Herting et al. 2015). The combination of at least two different sources of genetic and maturational influence into a unique descriptor of cortical volume may act as a confounding factor that hinders a clear interpretation in the context of cortical covariance based networks. This is particularly relevant in conditions such as autism and ADHD where differences in cortical thickness, cortical volume and surface area are highly heterogeneous (Wolosin et al. 2009; Ecker, Ginestet, et al. 2013).

Secondly, it is possible that in both publically available datasets, some participants might have been comorbid for the other condition (e.g., individuals in the ABIDE might have had comorbid ADHD, and *vice versa*). Although all individuals in these data-sets were diagnosed under the DSM-IV criteria, which does not allow this type of comorbidity, without the availability of more detailed diagnostic data, comorbidity cannot be ruled out completely. Yet the primary aim of this study was to investigate overlap between the two conditions. If the present results were due to the individuals that shared this comorbidity, this would still support a common underlying neural architecture. Nonetheless, future longitudinal studies need to disentangle this overlap more precisely and in relation to specific phenotypic overlap as well as the trajectory of topological changes over time.

In sum, we found convergence between autism and ADHD, where both conditions show stronger decrease in covariance with increased Euclidean distance between centroids compared to a neurotypical population. The two conditions also show divergence. Namely, there is less modular overlap between the two conditions then there is between each condition and the neurotypical group. The ADHD group also showed reduced cortical thickness and higher degree in hubs regions compared to the autism group. Lastly, the ADHD group also showed reduced wiring costs compared to the autism group. Future research investigating these patterns in functional and structural connectivity and relating findings to behavioural or phenotypic data will hopefully shed light on the convergent and divergent neural substrates of autism and ADHD. Our findings do support the notion that both developmental conditions involve a shift in network topology that might be characterized as favouring local over global patterns. Lastly, they highlight the value of taking an integrated approach across conditions.

## Acknowledgements

RB was funded by the Pinsent Darwin Trust, Medical Research Council UK and Cambridge Trust. SBC is supported by the National Institute for Health Research (NIHR) Collaboration for Leadership in Applied Health Research and Care East of England at Cambridgeshire and Peterborough NHS Foundation Trust. ETB is employed half-time by the University of Cambridge and half-time by GlaxoSmithKline (GSK); he holds stock in GSK. The views expressed are those of the author(s) and not necessarily those of the NHS, the NIHR, GSK or the Department of Health.

## References

Alexander-Bloch a., Raznahan a., Bullmore E, Giedd J. 2013. The Convergence of Maturational Change and Structural Covariance in Human Cortical Networks. J Neurosci. 33:2889–2899.

Alexander-Bloch A, Giedd JN, Bullmore E. 2013. Imaging structural co-variance between human brain regions. Nat Rev Neurosci. 14:322–336.

Almeida Montes LG, Prado Alcantara H, Martinez Garcia RB, De La Torre LB, Avila Acosta D, Duarte MG. 2013. Brain Cortical Thickness in ADHD: Age, Sex, and Clinical Correlations. J Atten Disord. 17:641–654.

American Psychiatric Association. 2013. Diagnostic and Statistical Manual of Mental Disorders. 5th ed.. Arlington, VA: American Psychiatric Press.

Assaf M, Jagannathan K, Calhoun V, Miller L, Stevens M, Sahl R, O’Boyle J, Schultz R, Pearlson G. 2010. Abnormal functional connectivity of default mode sub-networks in autism spectrum disorder patients. Neuroimage. 53:247–256.

Balardin JB, Comfort WE, Daly E, Murphy C, Andrews D, Murphy DGMGGM, Ecker C, Sato JR. 2015. Decreased centrality of cortical volume covariance networks in autism spectrum disorders. J Psychiatr Res. 69:142–149.

Belmonte MK, Allen G, Beckel-Mitchener A, Boulanger LM, Carper R a, Webb SJ. 2004. Autism and abnormal development of brain connectivity. J Neurosci. 24:9228–9231.

Benjamini Y, Hochberg Y. 1995. Controlling the False Discovery Rate: A Practical and Powerful Approach to Multiple Testing. J R Stat Soc Ser B. 57:289–300.

Brock J, Brown CC, Boucher J, Rippon G. 2002. The temporal binding deficit hypothesis of autism. Dev Psychopathol. 14:209–224.

Cai W, Chen T, Szegletes L, Supekar K, Menon V. 2015. Aberrant cross-brain network interaction in children with attention-deficit/hyperactivity disorder and its relation to attention deficits: a multi- and cross-site replication study. Biol Psychiatry. 1–11.

Cherkassky VL, Kana RK, Keller T a, Just MA. 2006. Functional connectivity in a baseline resting-state network in autism. Neuroreport. 17:1687–1690.

Courchesne E, Pierce K. 2005. Why the frontal cortex in autism might be talking only to itself: local over-connectivity but long-distance disconnection. Curr Opin Neurobiol. 15:225–230.

Desikan RS, Ségonne F, Fischl B, Quinn BT, Dickerson BC, Blacker D, Buckner RL, Dale AM, Maguire RP, Hyman BT, Albert MS, Killiany RJ. 2006. An automated labeling system for subdividing the human cerebral cortex on MRI scans into gyral based regions of interest. Neuroimage. 31:968–980.

Dougherty CC, Evans DW, Myers SM, Moore GJ, Michael AM. 2015. A Comparison of Structural Brain Imaging Findings in Autism Spectrum Disorder and Attention-Deficit Hyperactivity Disorder. Neuropsychol Rev. 25–43.

Ecker C, Ginestet C, Feng Y, Johnston P, Lombardo M V., Lai M-C, Suckling J, Palaniyappan L, Daly E, Murphy CM, Williams SC, Bullmore ET, Baron-Cohen S, Brammer M, Murphy DGM, MRC AIMS Consortium for the. 2013. Brain Surface Anatomy in Adults With Autism. JAMA Psychiatry. 70:59.

Ecker C, Ronan L, Feng Y, Daly E, Murphy C, Ginestet CE, Brammer M, Fletcher PC, Bullmore ET, Suckling J, Baron-Cohen S, Williams S, Loth E, Murphy DGM. 2013. Intrinsic gray-matter connectivity of the brain in adults with autism spectrum disorder. Proc Natl Acad Sci USA. 110:13222–13227.

Evans AC. 2013. Networks of anatomical covariance. Neuroimage. 80:489–504.

Fair DA, Posner J, Nagel BJ, Bathula D, Costa-Dias TG, Mills KL, Blythe MS, Giwa A, Schmitt CF, Joel T. 2010. Atypical Default Network Connectivity in Youth with ADHD. Biol Psychiatry. 68:1084–1091.

Gong G, He Y, Chen ZJ, Evans AC. 2012. Convergence and divergence of thickness correlations with diffusion connections across the human cerebral cortex. Neuroimage. 59:1239–1248.

Hardan AY, Libove RA, Keshavan MS, Melhem NM, Minshew NJ. 2009. A Preliminary Longitudinal Magnetic Resonance Imaging Study of Brain Volume and Cortical Thickness in Autism. Biol Psychiatry. 66:320–326.

Herting MM, Gautam P, Spielberg JM, Dahl RE, Sowell ER. 2015. A longitudinal study: changes in cortical thickness and surface area during pubertal maturation. PLoS One. 10:e0119774.

Just MA, Cherkassky VL, Keller T a, Kana RK, Minshew NJ. 2007. Functional and anatomical cortical underconnectivity in autism: evidence from an FMRI study of an executive function task and corpus callosum morphometry. Cereb Cortex. 17:951–961.

Just MA, Cherkassky VL, Keller TA, Minshew NJ. 2004. Cortical activation and synchronization during sentence comprehension in high-functioning autism: evidence of underconnectivity. Brain. 127:1811–1821.

Kern JK, Geier DA, King PG, Sykes LK, Mehta JA, Geier MR. 2015. Shared Brain Connectivity Issues, Symptoms, and Comorbidities in Autism Spectrum Disorder, Attention Deficit/Hyperactivity Disorder, and Tourette Syndrome. Brain Connect. 5:321–335.

Konrad K, Eickhoff SB. 2010. Is the ADHD brain wired differently? A review on structural and functional connectivity in attention deficit hyperactivity disorder. Hum Brain Mapp. 31:904–916.

Lee SH, Ripke S, Neale BM, Faraone S V, Purcell SM, Perlis RH, Mowry BJ, Thapar A, Goddard ME, Witte JS, Absher D, Agartz I, Akil H, Amin F, Andreassen OA, Anjorin A, Anney R, Anttila V, Arking DE, Asherson P, Azevedo MH, Backlund L, Badner JA, Bailey AJ, Banaschewski T, Barchas JD, Barnes MR, Barrett TB, Bass N, Battaglia A, Bauer M, Bayés M, Bellivier F, Bergen SE, Berrettini W, Betancur C, Bettecken T, Biederman J, Binder EB, Black DW, Blackwood DHR, Bloss CS, Boehnke M, Boomsma DI, Breen G, Breuer R, Bruggeman R, Cormican P, Buccola NG, Buitelaar JK, Bunney WE, Buxbaum JD, Byerley WF, Byrne EM, Caesar S, Cahn W, Cantor RM, Casas M, Chakravarti A, Chambert K, Choudhury K, Cichon S, Cloninger CR, Collier DA, Cook EH, Coon H, Cormand B, Corvin A, Coryell WH, Craig DW, Craig IW, Crosbie J, Cuccaro ML, Curtis D, Czamara D, Datta S, Dawson G, Day R, De Geus EJ, Degenhardt F, Djurovic S, Donohoe GJ, Doyle AE, Duan J, Dudbridge F, Duketis E, Ebstein RP, Edenberg HJ, Elia J, Ennis S, Etain B, Fanous A, Farmer AE, Ferrier IN, Flickinger M, Fombonne E, Foroud T, Frank J, Franke B, Fraser C, Freedman R, Freimer NB, Freitag CM, Friedl M, Frisén L, Gallagher L, Gejman P V, Georgieva L, Gershon ES, Geschwind DH, Giegling I, Gill M, Gordon SD, Gordon-Smith K, Green EK, Greenwood TA, Grice DE, Gross M, Grozeva D, Guan W, Gurling H, De Haan L, Haines JL, Hakonarson H, Hallmayer J, Hamilton SP, Hamshere ML, Hansen TF, Hartmann AM, Hautzinger M, Heath AC, Henders AK, Herms S, Hickie IB, Hipolito M, Hoefels S, Holmans PA, Holsboer F, Hoogendijk WJ, Hottenga J-J, Hultman CM, Hus V, Ingason A, Ising M, Jamain S, Jones EG, Jones I, Jones L, Tzeng J-Y, Kähler AK, Kahn RS, Kandaswamy R, Keller MC, Kennedy JL, Kenny E, Kent L, Kim Y, Kirov GK, Klauck SM, Klei L, Knowles JA, Kohli MA, Koller DL, Konte B, Korszun A, Krabbendam L, Krasucki R, Kuntsi J, Kwan P, Landén M, Långström N, Lathrop M, Lawrence J, Lawson WB, Leboyer M, Ledbetter DH, Lee PH, Lencz T, Lesch K-P, Levinson DF, Lewis CM, Li J, Lichtenstein P, Lieberman JA, Lin D-Y, Linszen DH, Liu C, Lohoff FW, Loo SK, Lord C, Lowe JK, Lucae S, MacIntyre DJ, Madden PAF, Maestrini E, Magnusson PKE, Mahon PB, Maier W,Malhotra AK, Mane SM, Martin CL, Martin NG, Mattheisen M, Matthews K, Mattingsdal M, McCarroll SA, McGhee KA, McGough JJ, McGrath PJ, McGuffin P, McInnis MG, McIntosh A, McKinney R, McLean AW, McMahon FJ, McMahon WM, McQuillin A,Medeiros H, Medland SE, Meier S, Melle I, Meng F, Meyer J, Middeldorp CM, Middleton L, Milanova V, Miranda A, Monaco AP, Montgomery GW, Moran JL, Moreno-De-Luca D, Morken G, Morris DW, Morrow EM, Moskvina V, Muglia P, Mühleisen TW, Muir WJ, Müller-Myhsok B, Murtha M, Myers RM, Myin-Germeys I, Neale MC, Nelson SF, Nievergelt CM, Nikolov I, Nimgaonkar V, Nolen WA, Nöthen MM, Nurnberger JI, Nwulia EA, Nyholt DR, O’Dushlaine C, Oades RD, Olincy A, Oliveira G, Olsen L, Ophoff RA, Osby U, Owen MJ, Palotie A, Parr JR, Paterson AD, Pato CN, Pato MT, Penninx BW, Pergadia ML, Pericak-Vance MA, Pickard BS, Pimm J, Piven J, Posthuma D, Potash JB, Poustka F, Propping P, Puri V, Quested DJ, Quinn EM, Ramos- Quiroga JA, Rasmussen HB, Raychaudhuri S, Rehnström K, Reif A, Ribasés M, Rice JP, Rietschel M, Roeder K, Roeyers H, Rossin L, Rothenberger A, Rouleau G, Ruderfer D, Rujescu D, Sanders AR, Sanders SJ, Santangelo SL, Sergeant JA, Schachar R, Schalling M, Schatzberg AF, Scheftner WA, Schellenberg GD, Scherer SW, Schork NJ, Schulze TG, Schumacher J, Schwarz M, Scolnick E, Scott LJ, Shi J, Shilling PD, Shyn SI, Silverman JM, Slager SL, Smalley SL, Smit JH, Smith EN, Sonuga-Barke EJS, St Clair D, State M, Steffens M, Steinhausen H-C, Strauss JS, Strohmaier J, Stroup TS, Sutcliffe JS, Szatmari P, Szelinger S, Thirumalai S, Thompson RC, Todorov AA, Tozzi F, Treutlein J, Uhr M, van den Oord EJCG, Van Grootheest G, Van Os J, Vicente AM, Vieland VJ, Vincent JB, Visscher PM, Walsh CA, Wassink TH, Watson SJ, Weissman MM, Werge T, Wienker TF, Wijsman EM, Willemsen G, Williams N, Willsey AJ, Witt SH, Xu W, Young AH, Yu TW, Zammit S, Zandi PP, Zhang P, Zitman FG, Zöllner S, Devlin B, Kelsoe JR, Sklar P, Daly MJ, O’Donovan MC, Craddock N, Sullivan PF, Smoller JW, Kendler KS, Wray NR. 2013. Genetic relationship between five psychiatric disorders estimated from genome-wide SNPs. Nat Genet. 45:984–994.

Leitner Y. 2014. The co-occurrence of autism and attention deficit hyperactivity disorder in children - what do we know? Front Hum Neurosci. 8:268.

Li X, Cao Q, Pu F, Li D, Fan Y, An L, Wang P, Wu Z, Sun L, Li S, Wang Y. 2015. Abnormalities of structural covariance networks in drug-naїve boys with attention deficit hyperactivity disorder. Psychiatry Res - Neuroimaging. 231:273–278.

Loth E, Spooren W, Ham LM, Isaac MB, Auriche-Benichou C, Banaschewski T, Baron-Cohen S, Broich K, Bölte S, Bourgeron T, Charman T, Collier D, de Andres-Trelles F, Durston S, Ecker C, Elferink A, Haberkamp M, Hemmings R, Johnson MH, Jones EJH, Khwaja OS, Lenton S, Mason L, Mantua V, Meyer-Lindenberg A, Lombardo M V., O’Dwyer L, Okamoto K, Pandina GJ, Pani L, Persico AM, Simonoff E, Tauscher-Wisniewski S, Llinares-Garcia J, Vamvakas S, Williams S, Buitelaar JK, Murphy DGM. 2015. Identification and validation of biomarkers for autism spectrum disorders. Nat Rev Drug Discov. 15:70–73.

Mostert JC, Shumskaya E, Mennes M, Onnink AMH, Hoogman M, Kan CC, Vasquez AA, Buitelaar J, Franke B, Norris DG, Arias A, Buitelaar J, Franke B, Norris DG. 2016. Characterising resting-state functional connectivity in a large sample of adults with ADHD. Prog Neuro-Psychopharmacology Biol Psychiatry. 67:82–91.

Panizzon MS, Fennema-Notestine C, Eyler LT, Jernigan TL, Prom-Wormley E, Neale M, Jacobson K, Lyons MJ, Grant MD, Franz CE, Xian H, Tsuang M, Fischl B, Seidman L, Dale A, Kremen WS. 2009. Distinct genetic influences on cortical surface area and cortical thickness. Cereb Cortex. 19:2728–2735.

Ray S, Miller M, Karalunas S, Robertson C, Grayson DS, Cary RP, Hawkey E, Painter JG, Kriz D, Fombonne E, Nigg JT, Fair DA. 2014. Structural and functional connectivity of the human brain in autism spectrum disorders and attention-deficit/hyperactivity disorder: A rich club-organization study. Hum Brain Mapp. 35:6032–6048.

Raznahan A, Lerch JP, Lee N, Greenstein D, Wallace GL, Stockman M, Clasen L, Shaw PW, Giedd JN. 2011. Patterns of coordinated anatomical change in human cortical development: a longitudinal neuroimaging study of maturational coupling. Neuron. 72:873–884.

Romero-garcia R, Atienza M, Clemmensen LH, Cantero JL. 2012. Effects of network resolution on topological properties of human neocortex. Neuroimage. 59:3522–3532.

Rommelse NNJ, Franke B, Geurts HM, Hartman CA, Buitelaar JK. 2010. Shared heritability of attention-deficit/hyperactivity disorder and autism spectrum disorder. Eur Child Adolesc Psychiatry. 19:281–295.

Rommelse NNJ, Geurts HM, Franke B, Buitelaar JK, Hartman CA. 2011. A review on cognitive and brain endophenotypes that may be common in autism spectrum disorder and attention-deficit/hyperactivity disorder and facilitate the search for pleiotropic genes. Neurosci Biobehav Rev. 35:1363–1396.

Rubenstein J, Merzenich M. 2003. Model of autism: increased ratio of excitation/inhibition in key neural systems. Genes, Brain Behav. 2:255–267.

Schmitt JE, Lenroot RK, Ordaz SE, Wallace GL, Lerch JP, Evans AC, Prom EC, Kendler KS, Neale MC, Giedd JN. 2009. Variance decomposition of MRI-based covariance maps using genetically informative samples and structural equation modeling. Neuroimage. 47:56–64.

Sharda M, Khundrakpam BS, Evans AC, Singh NC. 2014. Disruption of structural covariance networks for language in autism is modulated by verbal ability. Brain Struct Funct.

Shaw P, Eckstrand K, Sharp W, Blumenthal J, Lerch JP, Greenstein D, Clasen L, Evans A, Giedd J, Rapoport JL. 2007. Attention-deficit/hyperactivity disorder is characterized by a delay in cortical maturation. Proc Natl Acad Sci USA. 104:19649–19654.

Tomasi D, Volkow N. 2012. Abnormal functional connectivity in children with attention-deficit/hyperactivity disorder. Biol Psychiatry. 71:443–450.

van Wijk BCM, Stam CJ, Daffertshofer A. 2010. Comparing brain networks of different size and connectivity density using graph theory. PLoS One. 5.

Vissers ME, Cohen MX, Geurts HM. 2012. Brain connectivity and high functioning autism: a promising path of research that needs refined models, methodological convergence, and stronger behavioral links. Neurosci Biobehav Rev. 36:604–625.

Wolosin SM, Richardson ME, Hennessey JG, Denckla MB, Mostofsky SH. 2009. Abnormal cerebral cortex structure in children with ADHD. Hum Brain Mapp. 30:175–184.

Zielinski BA, Gennatas ED, Zhou J, Seeley WW. 2010. Network-level structural covariance in the developing brain. Proc Natl Acad Sci. 107:18191–18196.

